# Luciferase of the Japanese syllid polychaete *Odontosyllis umdecimdonta*

**DOI:** 10.1101/329631

**Authors:** Darrin T. Schultz, Alexey A. Kotlobay, Rustam Ziganshin, Artyom Bannikov, Nadezhda M. Markina, Tatiana V. Chepurnyh, Ekaterina S. Shakhova, Ksenia Palkina, Steven H.D. Haddock, Ilia V. Yampolsky, Yuichi Oba

## Abstract

1

*Odontosyllis undecimdonta* is a marine syllid polychaete that produces bright internal and exuded bioluminescence. Despite over fifty years of biochemical investigation into *Odontosyllis* bioluminescence, the light-emitting small molecule substrate and catalyzing luciferase protein have remained a mystery. Here we describe the discovery of a bioluminescent protein fraction from *O. undecimdonta,* the identification of the luciferase using peptide and RNA sequencing, and the *in vitro* reconstruction of the bioluminescence reaction using highly purified *O. undecimdonta* luciferin and recombinant luciferase. Lastly, we found no identifiably homologous proteins in publicly available datasets. This suggests that the syllid polychaetes contain an evolutionarily unique luciferase among all characterized luminous taxa.

**Highlights:**

- The polychaete *O. undecimdonta* uses a luciferin-luciferase bioluminescence system
- *O. undecimdonta* bioluminescence does not require additional cofactors
- The luciferase of the Japanese fireworm is 329 amino acids long
- Recombinant luciferase is not secreted when expressed in human cells
- Exogenous luciferin does not seem to penetrate cell membranes-only lysate luminesces
- The luciferase transcript is supported by full-length cDNA reads with 5’ and 3’ UTR

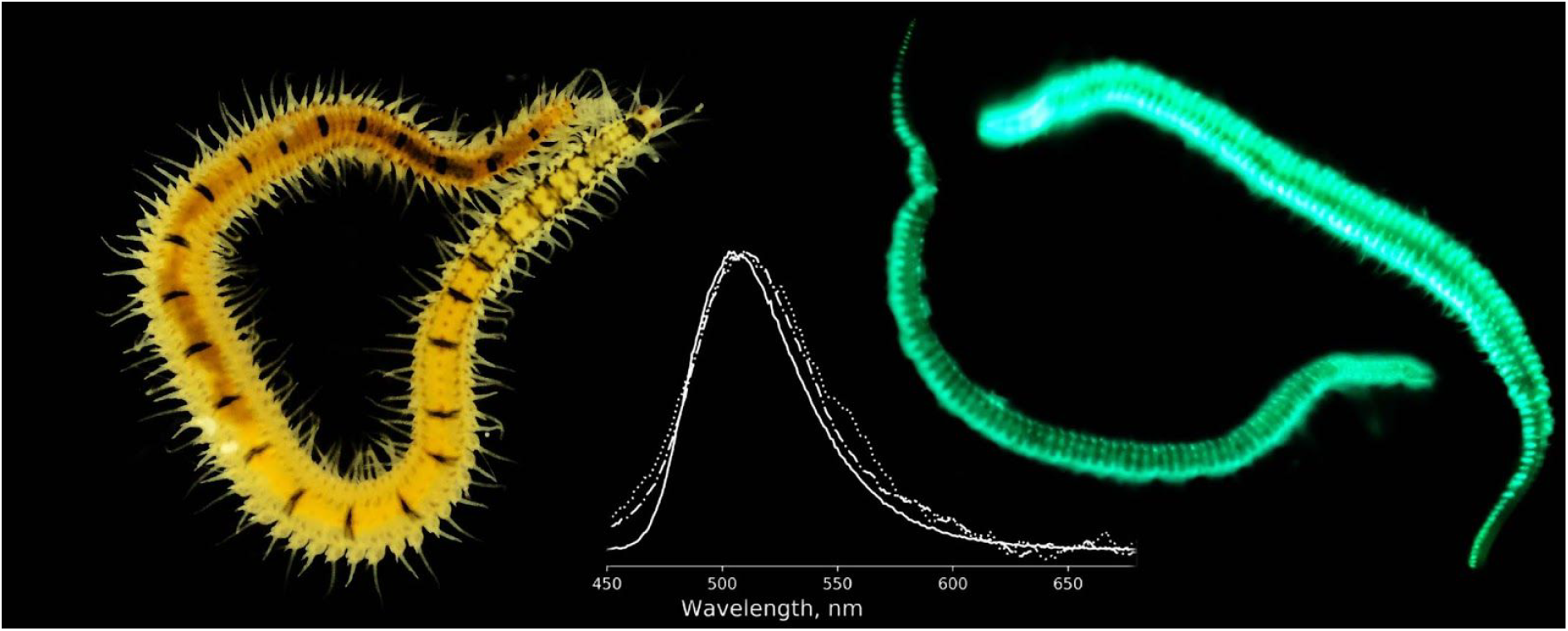

## 4 Introduction

*Odontosyllis* is a widely distributed genus of marine syllid polychaete worms that are noted for their striking bioluminescent courtship displays [1–5]. The bioluminescence (BL) of *Odontosyllis* is a luciferin-luciferase system [6], but the structure of the luciferin and the luciferase protein remain unknown despite several biochemical studies following the first in 1931 by Harvey [6–11]. More broadly, to date the enzyme sequences and luciferin structures remain a mystery for all polychaete species in the thirteen families containing luminous species [12].

Previous studies of the *Odontosyllis* bioluminescence system generated conflicting results regarding whether the system is a soluble oxygen-dependent luciferin-luciferase reaction [8,9], or is a photoprotein system in which the light-emitting small molecule substrate is covalently bound to the enzyme [11]. The above studies used a different *Odontosyllis* species, and the different colors of aqueous extracts identified from those species make it unclear whether there are multiple bioluminescent chemistries within *Odontosyllis*. However, both species have the same behavior of secreting luminescence during mating [1,4], so both species presumably share a homologous bioluminescent system.

*Odontosyllis undecimdonta* is a species found in Toyama Bay, Japan which engages in bioluminescent surface courtship displays around the first new moon in October [13]. Recently a protein-coding sequence from *O. undecimdonta* was patented that produces a recombinant protein with luminescence activity similar to that of crude worm extract mixed with crude luciferin isolate (WO2017155036A1). Here, we describe the identification, cloning, and characterization of the *O. undecimdonta* luciferase. In addition, our results suggest that the *O. undecimdonta* luminescence system is a luciferase-luciferin type without requisite cofactors, despite reports of magnesium ions as a necessary cofactor [14].

## 5 Materials and Methods

### 5.1 Specimen Collection

Professor S. Inoue provided lyophilized *O. undecimdonta* worms collected in 1993 to develop the protein purification strategy [15]. The final specimens used in this study for protein purification, MS transcript identification, and nucleic acid purification were collected on October 06, 2016 in Toyama Prefecture Japan, Namerikawa City. At dusk, *Odontosyllis* worms were attracted to a handheld light at the surface and collected with a hand dip net. Worms were individually preserved in Invitrogen RNAlater or lyophilized for later analysis.

#### 5.1.1 DNA and RNA isolation

Methods for DNA and RNA isolation, as well as construction of the RNA-seq and genomic DNA libraries are as described in the Supplementary Information. Briefly, the *O. undecimdonta* transcriptome was assembled using 32,457,166 Illumina 2×150 read pairs and 343,752 Oxford Nanopore long reads using the Trinity assembler [16]. DNA from a single *O. undecimdonta* specimen was used to prepare both a 10X Genomics chromium library [17] and a PCR-free library. All sequencing reads are available to download from the European Nucleotide Archive under project PRJEB26709. Individual luciferase transcripts are available at NCBI accession numbers MH350412 and MH350413.

### 5.2 Protein extraction from biomaterial

Five ml of phosphate buffer (5 mM sodium phosphate buffer, pH 7.4) was added to 150 mg of lyophilized worms. Then this mixture was dropped in to the liquid nitrogen, using a 1 ml pipette, to create small drops of frozen material. These small ice drops were ground in a mortar. Frozen powder was added to 10 ml of phosphate buffer (5 mM sodium phosphate buffer, pH 7.4) and incubated 40 min on an ice bath with stirring. After incubation this solution was centrifuged at 40000 g (4° C) for 40 min. The supernatant, containing luciferase, was then collected and used for further purification by anion exchange chromatography.

### 5.3 Protein purification

#### 5.3.1 Anion exchange chromatography of water extract

An extract of *O. undecimdonta* was applied to a DEAE Sepharose (GE Healthcare, Uppsala, Sweden) HiTrap Fast Flow column (1.6 ×; 2.5 cm), equilibrated and washed with 5 mM sodium phosphate buffer, pH 7.4 at rate of 5 ml/min. The elution was done by linear NaCl gradient from 0 to 0.4 M (80 ml) and 5 ml fractions were collected. To minimize bioluminescent reactions, the solvent, fractions and column were maintained at 4° C. Automatic fraction collection and solvent application was controlled with an Akta Prime chromatography system (GE Healthcare, Uppsala, Sweden). After elution, fractions containing luciferase and luciferin were detected by pairwise mixing all possible fraction combinations. The reaction was monitored with a custom-made luminometer Oberon-K (Krasnoyarsk, Russia).

#### 5.3.2 Ultrafiltration and concentration

To discard additional proteins from the luciferase-containing fractions the ultrafiltration procedure was used. First, the active fraction was filtered on a 50 kDa Amicon^®^ Ultra centrifugal filter unit (Merck Millipore, Germany). BL activity was measured for the concentrated retentate and the permeate. We found that only the permeate was bioluminescent. The bioluminescent permeate was then concentrated on 30 kDa Amicon^®^ Ultra centrifugal filter unit (Merck Millipore, Germany). The resulting retentate possessed BL activity while the permeate did not. Thus this concentrated luciferase sample was used for size exclusion chromatography.

#### 5.3.3 Size exclusion chromatography

The bioluminescent retentate from ultrafiltration was applied to a Superdex 200 column (Phenomenex, USA) on a Shimadzu chromatography system (Shimadzu, Japan). The loaded column was washed with 5 mM sodium phosphate buffer, 150 mM NaCl, pH 7.4 at rate of 0.4 ml/min. During separation 0.5 ml fractions were collected. The solvent, fractions, and column were maintained at 4° C. BL-active fractions were used in the subsequent gel electrophoresis experiments.

### 5.4 Denaturing polyacrylamide gel electrophoresis and amino acid sequence analysis

SDS-PAGE of the BL-active fractions was performed using a 10-25% gradient gel according to [18]. Gel staining was done according to the silver staining protocol from [19], or using a standard Coomassie G250 staining protocol. Protein bands were excised from the gel and subjected to in-gel trypsinolysis [20]. LC-MS was performed on the Ultimate 3000 Nano LC System (Thermo Fisher Scientific), connected to a Q Exactive HF mass spectrometer (Thermo Fisher Scientific). For data analysis, Mascot software (Matrix Science) with the *O. undecimdonta* transcriptome as a reference was used.

#### 5.4.1 Molecular cloning

Four Odontosyllis luciferase candidate genes were codon-optimized for expression in mammalian cells, domesticated for compatibility with MoClo assembly [21] and ordered from a commercial supplier (Twist Biosciences, USA) as linear dsDNA fragments. Molecular cloning is described in detail in the Supplementary Materials.

#### 5.4.2 Mammalian cell culture

HEK293NT cells were grown under standard conditions and transfected with FuGene 6 reagent (Promega, Fitchburg, WI, USA) in accordance to the manufacturer’s protocol. For more details see Supplementary materials.

### 5.5 In vitro bioluminescence assay

The reaction was monitored with a custom-made luminometer Oberon-K (Krasnoyarsk, Russia) at room temperature. For each measurement 100 μl of reaction mix (10 mM sodium phosphate buffer, 150 mM NaCl, pH 7.4, 2 μl of luciferase fraction, 2 μl of highly purified luciferin [22] were used. In experiments with mammalian cells lysates, the same amount of cells was used for each clone in each bioluminescence analysis to make results comparable.

The involvement of additional cofactors in the *O. undecimdonta* bioluminescence reaction was tested using an *in vitro* assay with only purified luciferase and highly purified luciferin. Since previous studies suggest the involvement of Mg^2+^ in the *Odontosyllis* bioluminescence reaction (optimum conc is 30 mM; [14]), we also testing the *in vitro* bioluminescence assay with 30 mM-60 mM Mg^2+^ with cell lysate.

### 5.6 Protein structure and homology analysis

HMMER and the BLAST suite were used to predict structural domains and interspecies homology of transcripts that produced bioluminescence [23–25]. We also used Phobius and SignalP to detect signal peptides and transmembrane domains of the same transcripts [26, 27]. Lastly, we used the I-TASSER server for structural prediction [28]. See the supplementary materials for a detailed description of the search for homologous sequences.

## 6 Results

The isolation and purification of *O. undecimdonta* luciferase required ion exchange chromatography, size exclusion chromatography, and ultrafiltration. (Fig. 1). The presence of luciferase in samples was controlled by an *in vitro* BL assay for all stages of purification. Several bands that corresponded to BL activity in the size exclusion chromatography fractions were identified by polyacrylamide gel electrophoresis (Fig. 1C). These bands were excised from gel and were identified by LC-MS.

**Fig. 1.**
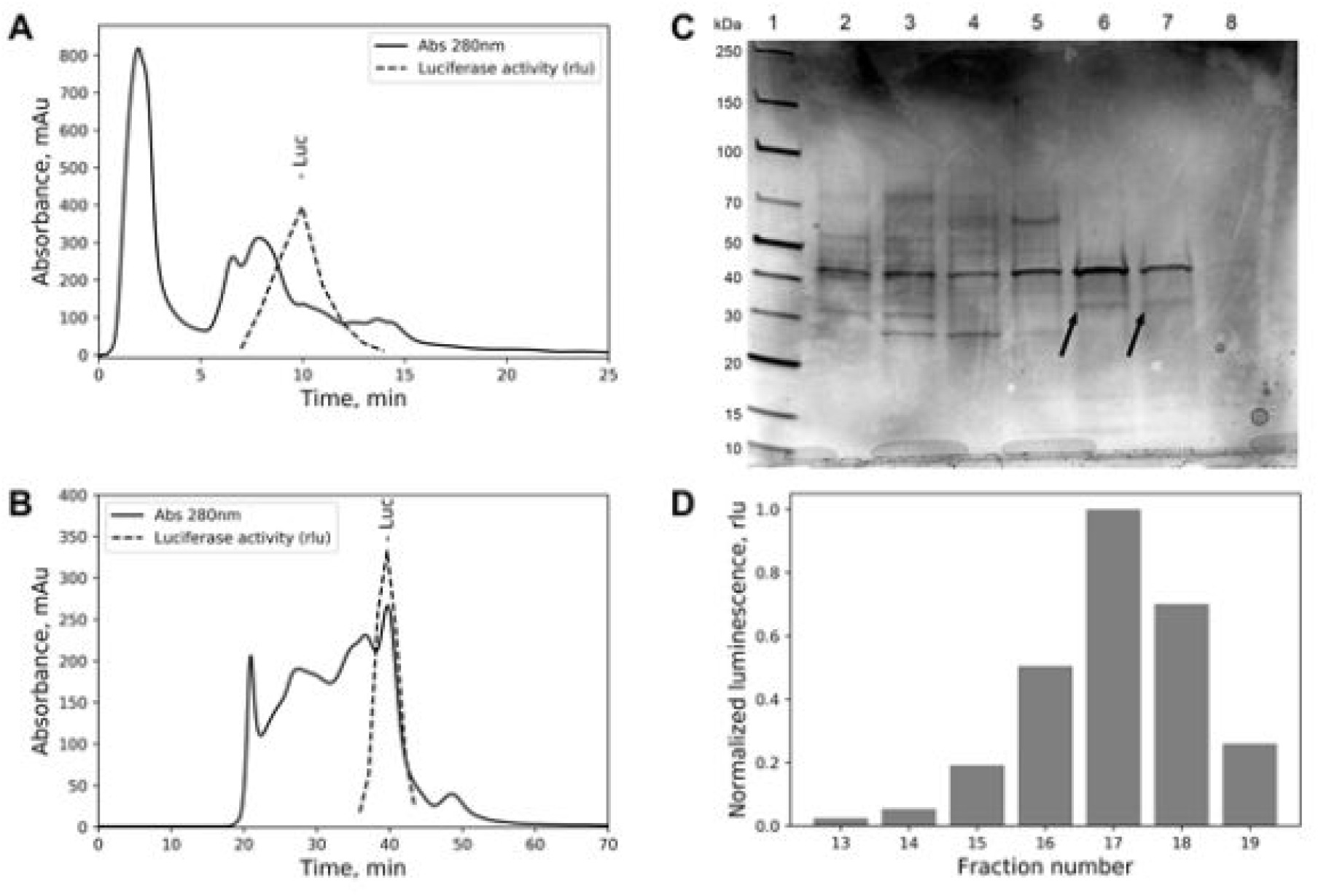
Purification and isolation of *O.undecimdonta* luciferase. **A** - Chromatographic profile of water extract from lyophilized *O. undecimdonta* worms anion-exchange chromatography on DEAE Sepharose HiTrap Fast Flow column. Solid line – 280 nm absorption signal. Activity profile of *Odontosyllis* luciferase shown as dashed line. **B** – Size exclusion chromatographic profile of anion-exchange chromatography concentrated luciferase fractions on Superdex 200 column. Solid line – 280 nm absorption signal. Activity profile of *O. undecimdonta* luciferase shown as dashed line. **C** - SDS-PAGE analysis of size exclusion chromatography fractions. Lanes: 1 - PageRuler protein ladder (Thermo Scientific), lanes from 2 to 8 - size exclusion chromatography fractions 13-19 respectively. Arrows shows protein bands excised from gel and analyzed by LC-MS. **D** – Normalized luminescence activity of size exclusion chromatography fractions, used for SDS-PAGE analysis.

The transcriptome assembled with Illumina paired-end reads and ONT 2D reads extracted with poretools “fwd” parameter yielded 256,027 transcripts and a median transcript length of 737 base pairs. Four transcripts were identified as potential luciferases (Fig. 2) based on coverage and quantity of MS matches. Three long transcripts c9g1i2 (990 bp), c9g1i3 (993 bp), c9g1i6 (990 bp) had c-terminal amino acid variation. Transcript c9g1i5 (711 bp) was homologous to the aforementioned three transcripts but lacked 118 n-terminal amino acids. These four transcripts were verified by presence of two ONT whole-cDNA reads that spanned from the 5’ UTR to the 3’UTR. Non-spliced mapping of an Illumina paired-end polyA RNA-seq library also confirmed that the longest of the four transcripts were expressed. The BLOSUM-alignment for the protein products of these four transcripts were identical at 92% of sites.

**Fig. 2.**
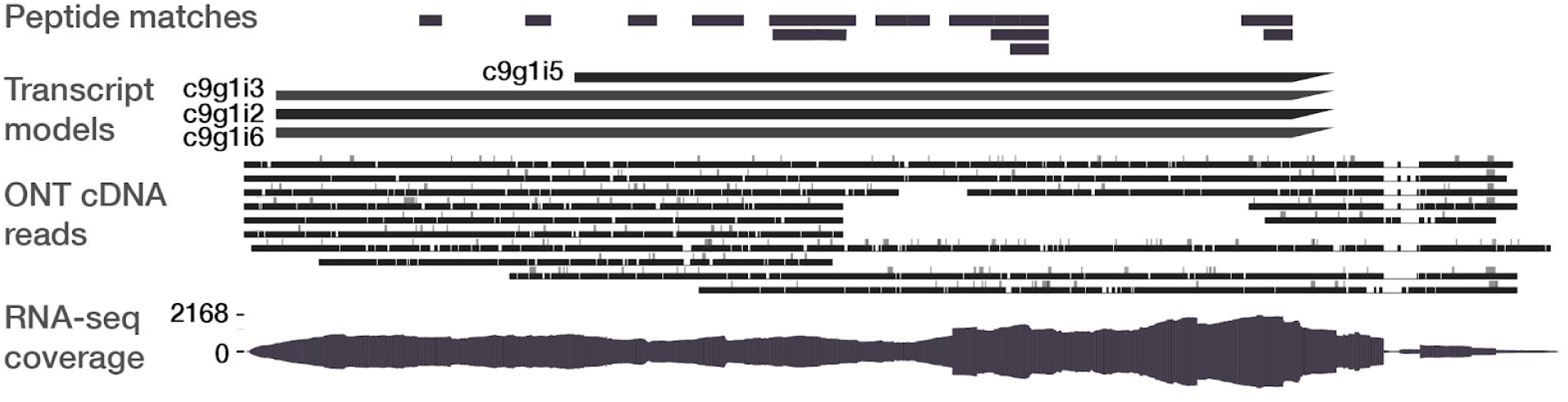
Supporting evidence for transcript models aligned to the c9g1i2 transcript, including 5’ and 3’ UTR. The *Peptide Matches* track shows unique peptide hits to any of the four transcript models that match by DNA and amino acid sequence similarity. All transcript models except c9g1i5 share the same structure, whereas c9g1i5 lacks 93 N-terminal amino acids. The *ONT cDNA Reads* track shows individual Oxford Nanopore 2D cDNA reads that align to the c9g1i2 transcript. Three reads span the complete 5’ UTR, transcript, and 3’ UTR of the long isoforms (c9g1i2, c9g1i3, c9g1i6), and four additional reads support the 5’ UTR of the long isoforms. The *RNA-seq coverage* track supports the 5’ and 3’ UTR of the long isoforms, despite the predictable 3’ bias inherent to polyA-selecting library preparation techniques.

All four candidate DNA sequences were synthesized as linear dsDNA fragments and cloned using MoClo technology. Then, mammalian cells were transfected by resulting constructs. Mammalian cell culture lysate from two of the above four candidates produced bioluminescence when assayed with purified luciferin (c9g1i2 and c9g1i6) (Fig. 3A). The bioluminescence spectra of positive clones were similar to that of native *O. undecimdonta* worms (Fig 3B). However, cell culture lysate from expressed transcripts c9g1i3 and c9g1i5 were not luminous. None of the non-lysed cell cultures produced luminescence when purified luciferin was applied.

**Fig. 3.**
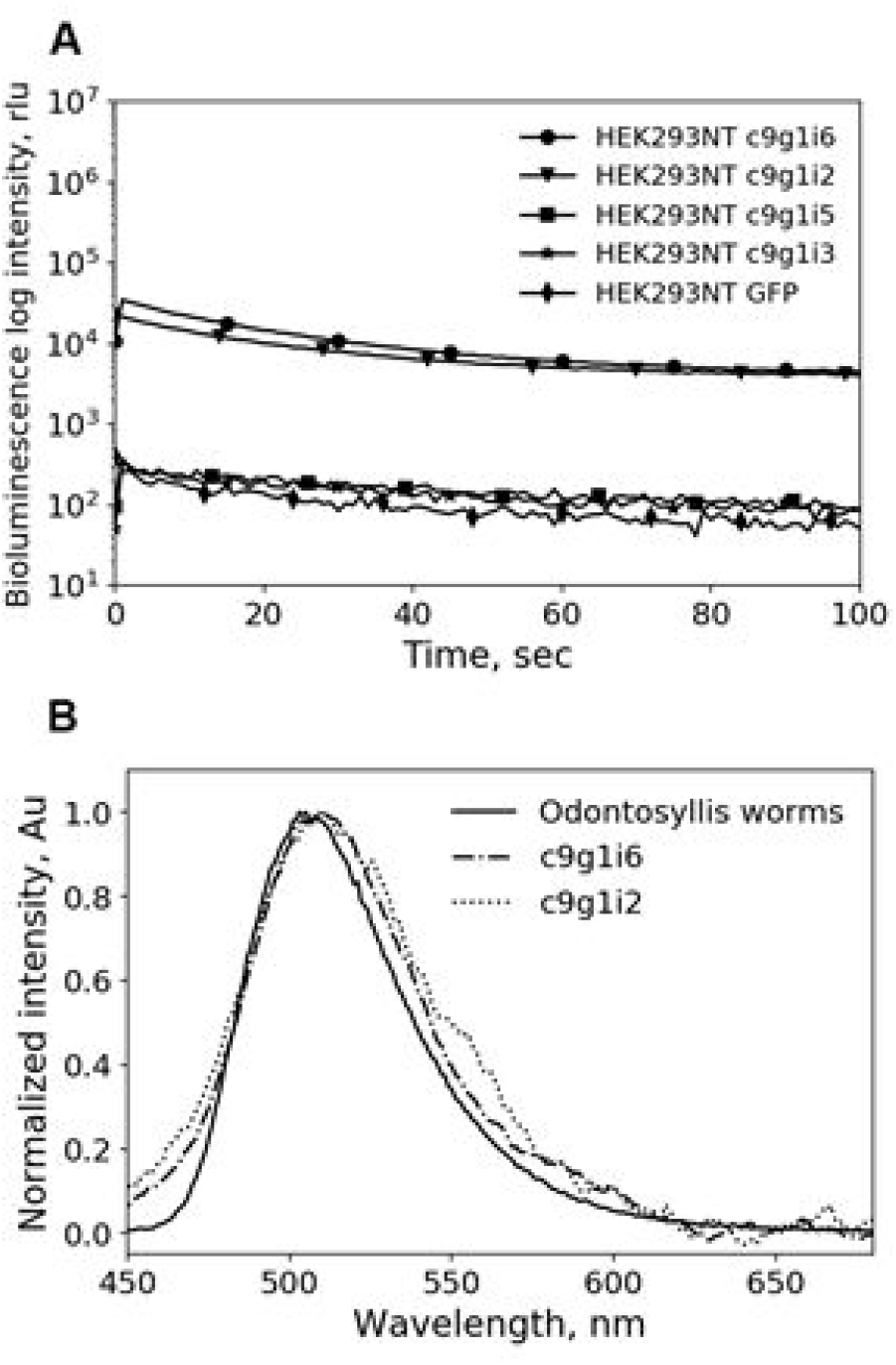
*O. umdecimdonta* luciferase properties. **A** - Kinetics of bioluminescence of transfected by *O. umdecimdonta* luciferase genes candidates HEK293NT cells lysates (log scale). For HEK293NT cells lysates, transfected by plasmids with c9g1i6, c9g1i2, c9g1i5, c9g1i3 - circle, triangle, square and asterisks markers were used respectively, and rhombus marker for GFP control. **B** – Normalized bioluminescence spectra of natural *O.undecimdonta* luciferase (solid line), c9g1i6 recombinant protein (dash-dotted line) and c9g1i2 recombinant protein (dotted line). The spectral maxima are near 510 nm.

**Fig. 4.**
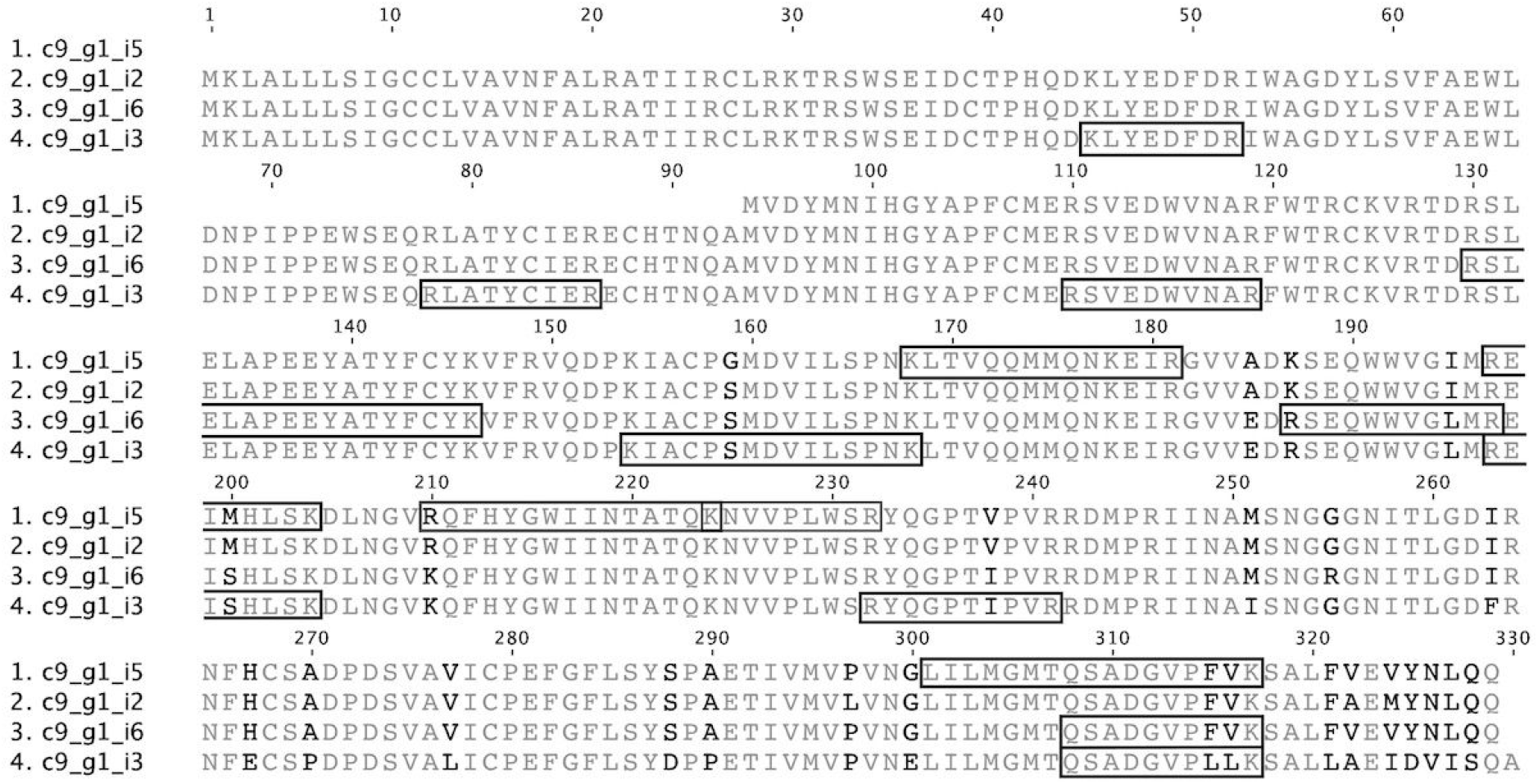
The amino acid alignment of the four luciferase transcripts. Black boxes surrounding the alignment indicate regions to which there were exact MS peptide matches. The four transcripts are on average 94% similar to one another. Transcript c9g1i5 lacks 93 N-terminal amino acids. All transcripts have a highly variable C-terminus.

**Fig. 5.**
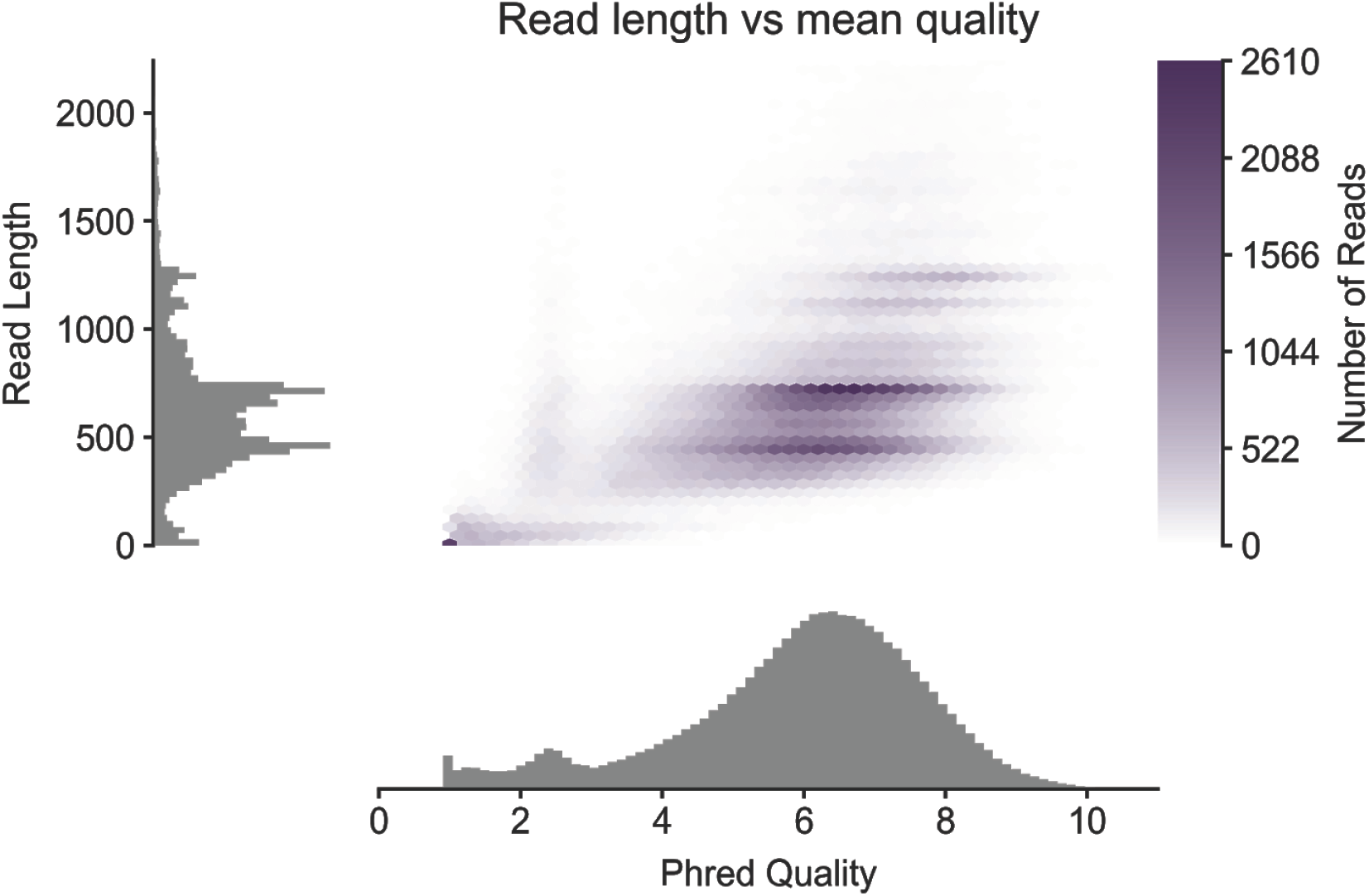
(SI) A marginplot of the basecalled 2D reads’ read lengths versus mean quality scores [23]. The average read quality is lower than what one would expect from a 2D sequencing run, however the read lengths indicate that many full-length transcripts were captured and sequenced as 2D reads.

The protein product of c9g1i2 is 329 amino acids long. The signal peptide prediction software Phobius had a posterior probability of 1 that the first 21 c-terminal peptides are a signal peptide. The SignalP service has a probability of 0.28 that the first 21 amino acids are signal peptides. The only HMMER and PHMMER results for this protein product were an insignificant match (E-value = 0.8) to a prokaryotic protein involved in mRNA production. I-TASSER protein structure and function prediction results found that nine of the top ten structural homologs to the protein product of c9g1i2 were adenosine deaminase/hydrolases. A tblastn search with the c9g1i2 protein product only found an insignificant match (E-value = 3.1) to a predicted gerbil transcription factor (sequence XM_021634012.1). A blastn search returned no significant matches. Blast searches against the assembled transcriptomes of publicly available polychaete RNA-seq read data also yielded no significant matches (SI results).

The mixture of purified *O. undecimdonta* luciferase and luciferin in TBS (50 mM Tris-HCl, 150 mM NaCl, pH 8.0) was luminescent, even in the absence of Mg^2+^ions. Increasing the Mg^2+^ concentration in the reaction buffer of recombinant luciferase cell lysate did not affect the yield of the bioluminescence reaction (data not shown).

## 7 Discussion

Given our lack of fresh specimens we opted to extract and purify the *Odontosyllis* luciferase directly from the lyophilized worms and successfully identified the luciferase gene using classic protein purification, luciferin purification, and recent whole-cDNA sequencing techniques. We then reconstructed native *Odontosyllis* bioluminescence *in vitro* using purified protein and highly purified luciferin [22] with no additional cofactors. Lastly, we verified the identity of the *Odontosyllis* luciferase gene by showing that recombinant protein and purified luciferin in cell-lysate is luminous, in which the luminescence spectra (λmax, near 510 nm) matches that of the *Odontosyllis in vivo* luminescence.

It is notable that using purified components in studying bioluminescence reactions is important to verify that off-target reactions are not the source of luminescence and to avoid erroneous interpretation of the results [14]. Given that the protein and luciferin purification products were luminous and that luminescence of recombinant luciferase cell lysates were not enhanced with Mg^2+^ the *O. undecimdonta* luciferase-luciferin reaction does not appear to require additional cofactors. It is also important to note that the recombinant protein is not secreted by eukaryotic cells and that the luminescence reaction only occurs when cells containing the recombinant luciferase are lysed. This suggests that the highly purified *O. undecimdonta* luciferin is not membrane-permeable, thus limiting the potential for applications of this luciferase in optogenetics or other cellular expression-based technology.

While the bioluminescence emitted during mating is well-characterized in *Odontosyllis* spp., the luciferase structure and the mechanism of the luciferin-luciferase reaction remains unclear. Despite this uncertainty, protein ortology searches using BLAST and HMMER show that syllid luciferase is unique both among sequenced polychaetes and other sequenced organisms in public databases. The lack of evidence for a conserved protein in the transcriptomes of other luminous polychaetes leaves open the theory that bioluminescence evolved more than three times in the annelids. In this conservative estimate, we only include the evolution of two unique bioluminescent systems for which either the structure of the luciferin, luciferase, or both have been determined (earthworms [29] and *Odontosyllis)*, plus at least one event for other annelids with uncharacterized bioluminescent systems.

Given that the structure of other polychaete luciferins is still unknown, this leaves the question of polychaete bioluminescence unanswered. Identification of the *O. umdecimdonta* luciferase sequence is the most important step to further characterization of this worm’s bioluminescent system and the screening of other purified polychaete luciferins for cross-reactivity.

## 8 Acknowledgements

We thank late Professor Shoji Inoue and Dr. Hisae Kakoi (Meijo University, Japan) for providing lyophilized *Odontosyllis* specimens, and Uozu Aquarium (Toyama, Japan) for help collecting *Odontosyllis* specimens.

## 9 Funding

This work was supported by the Russian Science Foundation grant 14-50-00131, the David and Lucile Packard Foundation, and the Monterey Bay Aquarium Research Institute. DTS was supported by the NSF GRFP DGE 1339067.

## Supplementary Material to

### Supplementary Methods

#### Genomic DNA isolation and sequencing

##### Genomic DNA Isolation

Genomic DNA of one *O. undecimdonta* specimen, OdonB, was isolated using the Omega Biotek E.Z.N.A. Mollusc DNA kit (product number D3373). A 30 mg sample of RNAlater-preserved tissue yielded 24 µg of DNA at 80 ng/µl in 300 µL. A 1 µl sample of OdonB DNA was imaged on a 1% agarose gel in a 150 V field for 45 minutes and was found to be larger than the 10 kbp ladder. We did not perform pulse field gel electrophoresis to image the size distribution of the DNA greater than 10 kbp.

##### Genome Library Prep

We prepared both a 10X Genomics chromium DNA library [1], as well as a PCR-free whole genome shotgun library. For the 10X Genomics chromium library preparation, we sent a sample of the DNA to the UC Davis DNA Technologies Core where they performed the library prep and a single lane of 2×150 PE sequencing on an Illumina HiSeq 4000.

To prepare a PCR-free whole genome shotgun library, we sheared 1.5 µg of DNA in 50 µl of 1xTE using a Bioruptor sonication device on setting HIGH, for 30 seconds ON/30 seconds OFF for 13 cycles until the DNA size distribution mode was 350bp. Every five shearing cycles we removed the DNA tube from the Bioruptor, vortexed it, and quickly spun down the contents. We used 1 µg of sheared DNA as input for the Illumina TruSeq DNA PCR-Free library prep kit. The final library concentration was 6.68 ng/µL and uses the TruSeq i7 index ACAGTG(A). This library was pooled and sent to the UC Davis DNA Technologies Core for 2×150 PE sequencing on an Illumina HiSeq 4000. The sequencing run produced 39,628,963 read pairs.

##### Genome Assembly

To assemble the genome, we used the 10X Genomics Supernova assembler v1.1.2. We opted to not use the PCR-free shotgun data to assemble the genome due to low predicted coverage, approximately 16x assuming a conservative guess of a 700 Mbp genome size. All computation was performed using Haddock lab computational resources at the Monterey Bay Aquarium Research Institute. We used the simple command “supernova run --id <runid> --fastqs <path to fastq>/ --localmem=500” to assemble the genome using 500 GB of memory and 96 cores [1]. This took approximately three days to complete.

#### RNA sequencing protocol and transcriptome assembly

##### RNA Isolation

Total RNA intended for an Illumina RNA-seq library was isolated using the Trizol protocol on an RNAlater-preserved specimen (OdonA) from the same collection location, date, and time as sample OdonB. The final RNA yield from a 40 mg OdonA Trizol extraction was 170 ng/µl quantified by nanodrop in 45 µl, or 7.65 µg.

We also isolated total RNA from another individual, OdonC, to use downstream for Oxford Nanopore cDNA full-length sequencing. We isolated total RNA from OdonC using the manufacturer’s recommended protocol. OdonC also has the same collection parameters as OdonB and OdonA. The final RNA concentration from approximately 30 mg of tissue was 14.2 µg from 142 ng/µl in 100 µl of ddH_2_O, quantified by qubit.

##### Illumina cDNA library prep and sequencing

A template-switching Illumina RNA-seq library from OdonB total RNA was prepared at Evrogen (Moscow, Russia) with a TruSeq Stranded mRNA Library Prep Kit v2 with the i7 index ACAGTG(A). The library was sequenced at the UC Davis DNA Technologies Core on an Illumina HiSeq 4000 2×150 PE run to a depth of 32,457,166 read pairs.

##### cDNA Synthesis for 2D ONT Sequencing

For cDNA sequencing on the Oxford Nanopore Technologies Minion, we first synthesized cDNA from sample OdonC. All primers used in the following protocol were adapted from [2]. To 50 ng OdonC total RNA in 8 µl, we added 2 µl of a modified SmartSeq2 Oligo dT primer (5’-/5Me-isodC/AAGCAGTGGTATCAACGCAGAGTACTTTTTTTTTTTTTTTTTTTTTTTTTTTTTTVN-3’) synthesized by IDT, 1 µL of 10 mM dNTPs. We mixed by vortexing and spun down briefly. We incubated this mixture at 65° C for five minutes and snap-cooled on a freezer block in ice. To this reaction we added 4 µl of 5x RT buffer, 1 µl of 100 mM DTT, 1 µl of RNaseOUT, and 2 µl of 10 mM strand-switching oligo (5’-AAGCAGTGGTATCAACGCAGAGTACATrGrGrG-3’). This mixture was mixed by vortexing and spun down briefly, then incubated for 2 minutes at 42° C on a thermal cycler. Then, 1 µl of SuperScript IV enzyme (200 U/µl) was added to the mixture and mixed with five 1 µl pipette strokes. The reverse transcription reaction was carried out as follows: 10 minutes at 50°C for RT, then 10 minutes at 42°C for strand switching, then 10 minutes at 80°C for heat inactivation.

To amplify the cDNA, we set up three PCR reactions using the above RT reaction as input: 5 µl of the RT reaction, 1.5 µl of 10 mM ISPCR primer (5′-AAGCAGTGGTATCAACGCAGAGT-3′), 18.5 µl NFW, and 25 µl of LongAmp Taq 2x Master Mix. Reaction contents were mixed by gentle inversion then centrifuged to remove bubbles. The PCR reaction conditions were: one cycle of 95°C for 10 seconds, fifteen cycles of 95°C for 15 seconds then 64°C for 15 seconds then 65°C for 500 seconds, and one cycle of 65°C for 10 minutes.

The resulting cDNA was visualized on an agarose gel and all three amplicons were pooled together.

##### Library Prep and 2D ONT sequencing

From the pooled OdonC cDNA 1 µg was used as input for the remainder of the standard SQK-LSK208 Oxford Nanopore Technologies 2D Strand switching cDNA sequencing protocol. The final library concentration after 2D adapter-ligated capture and prior to sequencing was 8.18 ng/µl. The final library mass loaded to the flow cell was 98.16 ng in 12 µl of library. The flow cell used was a model FLO-MIN106, and the flowcell ID was FAF06207. We used MinKNOW v1.3.30 to control the sequencing run.

The sequencing run produced 428,172 fast5 read files. We used Albacore v1.1.1 to perform 2D basecalling on the reads and poretools v0.6.0 to extract reads from the basecalled fast5 files [3].

##### Transcriptome Assembly

Adapters were trimmed from the Illumina RNA-seq reads using SeqPrep2 [4]. We then *de novo* assembled a transcriptome using Trinity v2.1.1 with the option --SS_lib_type FR for read directionality and the --long_reads option using all 2D reads extracted from the Albacore-basecalled Oxford Nanopore reads [5].

##### Read Mapping

Illumina RNA-seq reads were mapped to the transcripts with bwa mem [6]. Oxford Nanopore Technologies cDNA reads were mapped to the transcripts with the splice-aware minimap2 [7]. Peptide matches were extracted from source transcripts, then the small sequences were mapped to the reference transcript with bwa aln [8]. This is the information that is displayed in Figure 2 of the main text. This procedure allows some amino acid mismatch when matching mass spectrometry hits to a DNA sequence. This provides the advantage of finding signal when population-level amino acid diversity is high.

##### Mammalian cell culture

HEK293NT were maintained in DMEM supplemented with 10% FBS and 1×; Penicillin/Streptomycin (“fullDMEM”) for all growth and passaging steps unless otherwise noted. For continuous culture, the cells were grown to approximately 70–80% confluency and then split 1:12 to be ready to be split again 3 days later. To split cells from a 25 cm^2^ flask the culture medium was gently removed, 1 ml PBS without Mg/Ca was added to cover the surface of the cells. PBS was removed and 1 ml 0.025 % Trypsin in 6 mM EDTA was added to the side of the flask, not directly onto the cells. Solution was spread over the cells by gently “rocking” the flask several times. The flask was incubated at 37° C for 1–2 min. Then flask was rocked to completely dislodge the cells. After gently pipetting 80 µl of cell suspension was transferred to the new flask supplied with 5 ml of fullDMEM.

For preparing cells for transfection, 40 µl of cell suspension described above was transferred to 2 cm cell culture dish supplied with 2 ml of fullDMEM. After 24 hours incubation at 37° C with 5% CO_2_ cells were transfected with FuGene 6 reagent (Promega, Fitchburg, WI, USA) in accordance to the manufacturer’s protocol.

##### Molecular cloning

All cloning was performed by Golden Gate assembly. The synthesized genes were cloned into MoClo Level 0-SP vector from MoClo kit plasmid pICH41258. Level 1 eukaryotic expression plasmids were assembled into MoClo kit plasmid pICH47742 as a backbone, and the following parts were cloned in Level 0 vectors: CMV promoter, luciferase candidate gene, stop-codon containing DNA part and SV40poyA terminator. Prokaryotic expression plasmids were assembled with pCOOFY plasmid (T7 promoter) as a backbone and luciferase candidate gene as a single insert. A vector containing GFP was used as a positive control for cloning and expression.

##### Sequences for Molecular Cloning

Below are the DNA sequences of the dsDNA transcripts ordered from Twist for cloning into MoClo Level 0-SP.

>Odontosyllis luciferase candidate #1 (short ORF from DN46871_c9_g1_i5), seq to order at Twist

GAAGACaaaATGGTGGACTATATGAATATTCATGGATATGCCCCTTTTTGCATGGAACGTAGTGTTgagGACTGGGTGAATGCTCGTTTCTGGACTCGTTGTAAGGTTCGTACTGATCGTAGTTTAGAACTGGCACCTGAAGAATATGCCACCTACTTTTGTTATAAGGTGTTTCGTGTAcaaGATCCTAAAATTGCTTGTCCAGGAATGGATGTGATCCTTTCACCTAACAAACTGACTGTACAACAAATGATGCAGAATAAGGAAATTCGTGGAGTTGTAGCAGATAAATCTGAGCAATGGTGGGTTGGAATTATGCGTGAAATTATGCATCTGTCTAAGGACTTGAATGGTGTTCGTCAATTCCATTATGGATGGATCATCAACACAGCTACACAAAAGAATGTGGTTCCTTTGTGGTCACGTTATcaaGGACCTACTGTTCCAGTACGTCGTGACATGCCTCGTATCATTAATGCCATGTCTAATGGCGGAGGAAACATCACCTTGGGAGATATTCGTAATTTCCACTGCTCTGCTGATCCAGACAGTGTTGCTGTCATCTGCCCTGAGTTTGGTTTCTTGTCCTATtcacccGCTGAAACTATCGTTATGGTTCCAGTAAATGGATTAATCCTGATGGGAATGACACAATCTGCAGATGGAGTACCCTTCGTAAAATCTGCCCTGTTTGTTGAGGTGTATAACTTGCAACAGtcaggtaaGTCTTC

>Odontosyllis luciferase candidate #2 (long ORF from DN46871_c9_g1_i2), seq to order at Twist

GAAGACaaaATGAAGTTAGCACTGTTATTAAGTATTGGATGTTGCCTGGTTGCCGTGAACTTTGCTTTACGTGCTACTATCATTCGTtgcCTTCGTAAAACTCGTAGTTGGTCAGAAATTGATTGTACACCACATCAGGACAAGCTGTATGAGGACTTTGACCGTATCTGGGCCGGAGATTACCTGTCAGTATTTGCTGAATGGTTAGATAATCCCATCCCCCCAGAGTGGTCTGAGCAACGTCTGGCCACATACTGCATTGAGCGTGAATGTCACACTAATCAAGCTATGGTTGACTATATGAATATCCATGGATATGCCCCTTTTTGCATGGAACGTAGTGTTgagGACTGGGTGAATGCTCGTTTCTGGACTCGTTGTAAGGTTCGTACTGACCGTAGTTTAGAACTGGCACCTGAAGAATATGCCACCTACTTTTGTTATAAGGTGTTTCGTGTAcaaGATCCTAAAATTGCTtgcCCCTCCATGGATGTGATCCTTTCACCTAACAAACTGACTGTACAACAAATGATGCAAAATAAGGAAATTCGTGGAGTTGTAGCAGATAAATCTGAGCAATGGTGGGTTGGAATTATGCGTGAAATCATGCATCTGTCTAAGGACTTGAATGGTGTTCGTCAATTCCATTATGGATGGATCATTAACACAGCTACACAAAAGAATGTGGTTCCTTTGTGGTCACGTTATcaaGGACCTACTGTTCCAGTACGTCGTGACATGCCTCGTATCATTAATGCCATGTCTAATGGCGGAGGAAACATCACCTTGGGAGATATTCGTAATTTCCACTGCtccGCTGATCCAGACAGTGTTGCTGTCATCTGCCCTGAGTTTGGTTTCTTGTCCTATAGTcctGCTGAAACTATTGTTATGGTTCTTGTAAATGGATTAATCCTGATGGGAATGACACAATCTGCAGATGGTGTACCCTTCGTAAAATCTGCACTGTTTGCTGAGATGTATAACCTTCAACAGtcaggtaaGTCTTC

> Odontosyllis luciferase candidate #3 (long ORF from DN46871_c9_g1_i3), seq to order at Twist

GAAGACaaaATGAAGTTAGCACTGTTATTATCTATTGGATGTTGCCTGGTTGCCGTGAACTTTGCTTTACGTGCTACTATCATCCGTtgcCTTCGTAAAACTCGTAGTTGGTCAGAAATTGATTGTACACCACATCAGGACAAGCTGTATGAGGACTTTGACCGTATCTGGGCCGGAGATTACCTGTCAGTATTTGCTGAATGGTTAGATAATCCCATCCCCCCAGAGTGGTCTGAGCAACGTCTGGCCACATACTGCATTGAGCGTGAATGTCACACTAATCAAGCTATGGTTGACTATATGAATATCCATGGATATGCCCCTTTTTGCATGGAACGTAGTGTTgagGACTGGGTGAATGCTCGTTTCTGGACTCGTTGTAAGGTTCGTACTGACCGTAGTTTAGAACTGGCACCTGAAGAATATGCCACCTACTTTTGTTATAAGGTGTTTCGTGTAcaaGATCCTAAAATTGCTtgcCCCTCAATGGATGTGATCCTTTCACCTAACAAACTGACTGTACAACAAATGATGCAAAATAAGGAAATCCGTGGAGTTGTAGAGGATCGTTCTGAGCAATGGTGGGTTGGACTGATGCGTGAAATTAGTCATCTGTCTAAGGACTTGAATGGTGTGAAACAATTCCATTATGGATGGATCATCAACACAGCTACACAAAAGAATGTGGTTCCTTTGTGGTCACGTTATCAGGGTCCTACTATTCCAGTACGTCGTGACATGCCTCGTATCATTAATGCCATTTCTAATGGAGGAGGAAACATCACCTTGGGAGATTTTCGTAATTTTGAATGCTCACCTGATCCAGATAGTGTTGCTCTGATCTGCCCTGAGTTTGGTTTCTTGTCCTATGATCCCCCTGAAACTATTGTAATGGTGCCAGTAAATGAATTAATCCTGATGGGAATGACACAATCTGCTGATGGAGTACCTTTGTTGAAGTCTGCCCTTTTAGCTGAGATTGATGTCATTTCCCAAGCTtcaggtaaGTCTTC

>Odontosyllis luciferase candidate #4 (long ORF from DN46871_c9_g1_i6), seq to order at Twist

GAAGACaaaATGAAGTTAGCACTGTTACTTTCTATTGGATGTTGCCTGGTTGCCGTGAACTTTGCTTTACGTGCTACTATCATTCGTtgcCTTCGTAAAACTCGTAGTTGGTCAGAAATCGATTGTACACCACATCAGGACaaaCTTTATGAGGACTTTGACCGTATCTGGGCCGGAGATTACCTGTCAGTATTTGCTGAATGGTTAGATAATCCCATCCCCCCAGAGTGGTCTGAGCAACGTCTGGCCACATACTGCATTGAGCGTGAATGTCACACTAATCAAGCTATGGTTGACTATATGAATATCCATGGATATGCCCCTTTTTGCATGGAACGTAGTGTTgagGACTGGGTGAATGCTCGTTTCTGGACTCGTTGTAAGGTTCGTACTGACCGTAGTTTAGAACTGGCACCTGAAGAATATGCCACCTACTTTTGTTATAAGGTGTTTCGTGTAcaaGATCCTAAAATCGCTtgcCCCTCCATGGATGTGATCCTTTCACCTAACAAACTGACTGTACAACAAATGATGCAAAATAAGGAAATTCGTGGAGTTGTAGAGGATCGTTCTGAGCAATGGTGGGTTGGATTGATGCGTGAAATCTCCCATCTGTCTAAGGACTTGAATGGTGTGAAACAATTCCATTATGGATGGATCATCAACACAGCTACACAAAAGAATGTGGTTCCTTTGTGGTCACGTTATCAGGGTCCTACTATTCCAGTACGTCGTGACATGCCTCGTATCATTAATGCCATGTCTAATGGCCGTGGAAACATCACCTTGGGAGATATTCGTAATTTCCACTGCtctGCTGATCCAGACAGTGTTGCTGTCATCTGCCCTGAGTTTGGTTTCTTGTCCTATTCAcccGCTGAAACTATCGTTATGGTTCCAGTAAATGGATTAATCCTGATGGGAATGACACAATCTGCAGATGGAGTACCCTTCGTAAAATCTGCCCTTTTTGTTGAGGTGTATAACCTGCAACAGtcaggtaaGTCTTC

t

>Odontosyllis luciferase candidate #4 (long ORF from DN46871_c9_g1_i6)

ATGAAGTTAGCACTGTTACTTTCTATTGGATGTTGCCTGGTTGCCGTGAACTTTGCTTTACGTGCTACTATCATTCGTtgcCTTCGTAAAACTCGTAGTTGGTCAGAAATCGATTGTACACCACATCAGGACaaaCTTTATGAGGACTTTGACCGTATCTGGGCCGGAGATTACCTGTCAGTATTTGCTGAATGGTTAGATAATCCCATCCCCCCAGAGTGGTCTGAGCAACGTCTGGCCACATACTGCATTGAGCGTGAATGTCACACTAATCAAGCTATGGTTGACTATATGAATATCCATGGATATGCCCCTTTTTGCATGGAACGTAGTGTTgagGACTGGGTGAATGCTCGTTTCTGGACTCGTTGTAAGGTTCGTACTGACCGTAGTTTAGAACTGGCACCTGAAGAATATGCCACCTACTTTTGTTATAAGGTGTTTCGTGTAcaaGATCCTAAAATCGCTtgcCCCTCCATGGATGTGATCCTTTCACCTAACAAACTGACTGTACAACAAATGATGCAAAATAAGGAAATTCGTGGAGTTGTAGAGGATCGTTCTGAGCAATGGTGGGTTGGATTGATGCGTGAAATCTCCCATCTGTCTAAGGACTTGAATGGTGTGAAACAATTCCATTATGGATGGATCATCAACACAGCTACACAAAAGAATGTGGTTCCTTTGTGGTCACGTTATCAGGGTCCTACTATTCCAGTACGTCGTGACATGCCTCGTATCATTAATGCCATGTCTAATGGCCGTGGAAACATCACCTTGGGAGATATTCGTAATTTCCACTGCtctGCTGATCCAGACAGTGTTGCTGTCATCTGCCCTGAGTTTGGTTTCTTGTCCTATTCAcccGCTGAAACTATCGTTATGGTTCCAGTAAATGGATTAATCCTGATGGGAATGACACAATCTGCAGATGGAGTACCCTTCGTAAAATCTGCCCTTTTTGTTGAGGTGTATAACCTGCAACAGtcaggt

#### Methods for BLAST search for luciferase homolog search

Trinity to assemble transcriptomes from publicly available polychaete RNA-seq data of the following species: Amphinomidae (*Pareurythoe californica* SRR1926090 [9]), Chaetopteridae (*Chaetopterus sp.* SRR1646443 [10], *Chaetopterus variopedatus* SRR5590967, *Mesochaetopterus minutus* SRR1925760 [9], *Phyllochaetopterus sp.* SRR1257898 [11], *Spiochaetopterus sp.* SRR1224605 [11]), Eunicida (*Eunice pennata* SRR2040479, *Eunice torquata SRR2005375 [9]*), Cirratulidae (*Cirratulus cirratus* SRR5590966, *Cirratulus spectabilis* SRR3574861 [12]), Flabelligeridae (*Flabelligera mundata* SRR3574613 [12]), Acrocirridae (*Macrochaeta clavicornis* SRR1221445 [11]), Phyllodocidae (*Phyllodoce medipapillata* SRR2016923 [9]), Polynoidae (*Harmothoe extenuata* SRR1237766 [11], *Harmothoe imbricata* SRR2005364 [9] and SRR4841788 [13]), Syllidae (*Syllis sp.* SRR1224604 [11]), and Tomopteridae (*Tomopteris helgolandica* SRR1237767 [11]).

To search for luciferase homologs, we assembled transcriptomes of polychaetes using publicly available data. To do this, we downloaded the forward and reverse read fastq.gz files from the European Nucleotide Archive. The confirmed luminous species included in this analysis were *Chaetopterus variopedatus* [14], *Harmothoe extenuata* [15], *Harmothoe imbricata* [16], and *Tomopteris helgolandica* [17]. All other species mentioned above may be luminous, with the exception of: *Eunice spp., Pareurythoe californica, Phyllodoce medipapillata* [18]. Below is a list of species included in this analysis.

- Amphinomidae
  - Pareurythoe californica SRR1926090 [9]
- Chaetopteridae
  - Chaetopterus sp. SRR1646443 [10]
  - Chaetopterus variopedatus SRR5590967
  - Mesochaetopterus minutus SRR1925760 [9]
  - Phyllochaetopterus sp. SRR1257898 [11]
  - Spiochaetopterus sp. SRR1224605 [11]
- Eunicida
  - Eunice pennata SRR2040479
  - Eunice torquata SRR2005375 [9]
- Cirratulidae
  - Cirratulus cirratus SRR5590966
  - Cirratulus spectabilis SRR3574861 [12]
  - Flabelligeridae (Flabelligera mundata SRR3574613 [12]
- Acrocirridae
  - Macrochaeta clavicornis SRR1221445 [11]
- Phyllodocidae
  - Phyllodoce medipapillata SRR2016923 [9]
- Polynoidae
  - Harmothoe extenuata SRR1237766 [11]
  - Harmothoe imbricata SRR2005364 [9]
  - Harmothoe imbricata scale SRR4841788 [13]
- Syllidae
  - Syllis sp. SRR1224604 [11]
- Tomopteridae
  - Tomopteris helgolandica SRR1237767 [11]

Reads were trimmed using Trimmomatic paired-end version 0.35 with the options ILLUMINACLIP:TruSeq3-PE-2.fa:2:30:10 LEADING:3 TRAILING:3 SLIDINGWINDOW:4:15 MINLEN:36 [19]. Trimmed reads were used to assemble each transcriptome using Trinity version 2.6.6 with default parameters [5]. The resulting transcriptome nucleotide fasta file was used in subsequent tblastn searches [20]. The nucleotide transcriptome was translated into longest complete ORFs using transdecoder version 5.3.0 [21]. This protein fasta file was used for blastp searches [20].

To search for homologs of the putative *O. undecimdonta* luciferase, we used the protein product of transcript c9g1i2 as a query against individual translated polychaete transcriptomes. For each blastp search, we limited the search to the top hit using -max_target_seqs 1. The name of the top hit, percent identity of the blastp alignment, the length of the alignment, and the E-value of the blastp hit are reported in column *‘c9g1i2 queried against transcriptomes’* of table 1 (SI). To determine if the best blastp match in each polychaete transcriptome had a match to a known non-luciferase protein, the top blastp hit was used as a query in a blastp search against the nr database. The results of these blast searches are reported in the column *“top hit from ‘c9g1i2 queried against transcriptomes’ queried against nr”* of table 1 (SI).

**Table 1.**
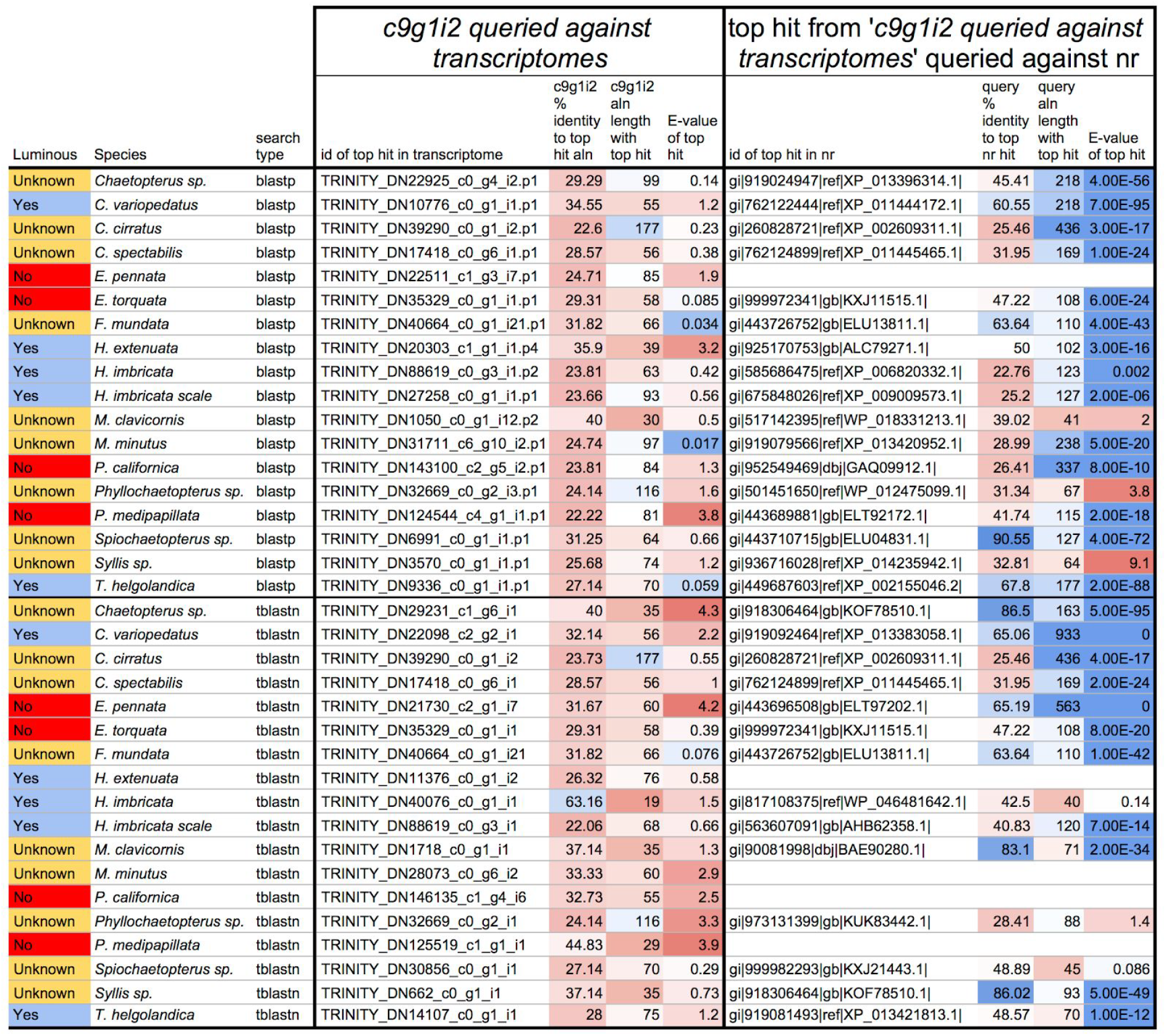
(SI) Blast results from individual searches against polychaete transcriptomes. The top half of the figure shows results from using c9g1i2 as the blastp query against translated polychaete transcriptomes, the bottom half shows results of using c9g1i2 as a tblastn query against untranslated polychaete transcriptomes. All blast result cell colors are colored on a scale of red to white to blue, wherein red indicates a dissimilar blast result while blue indicates a similar blast result. The ‘% identity to top hit aln’ columns show the percent identity match for the blast alignments. The ‘aln length with top hit’ columns show the length of the blast matches.

To include the possibility that a homologous sequence was not translated by the Transdecoder software, we performed another search using the c9g1i2 protein sequence as a tblastn query against individual polychaete nucleotide transcriptomes. The standard translation table for each polychaete genome was used when performing the tblastn search (-db_gencode 1). As above, the blast results for this search are listed in the *‘c9g1i2 queried against transcriptomes’* column of table 1 (SI). The top tblastn hit was used as a blastx query against the nr database. The standard translation table was used for each query (-query_gencode 1). As above, the results of the blastx search are listed in column *“top hit from ‘c9g1i2 queried against transcriptomes’ queried against nr”* of table 1 (SI). Blast searches with no results are listed as a blank line in table 1 (SI).

This script is available on github and is archived on zenodo [22].

### Supplementary Results

#### ONT cDNA sequencing results

With poretools we extracted the following quantities and types of reads: 421,040 forward reads, 343,752 2D reads, and 23,566 2D high quality reads. The total basecalled and extracted data yield was 328.4 million basepairs. The read length N50 was 836 basepairs. A hex-bin density plot and marginal density plot for the untrimmed reads are visualized in Figure 6.

#### Genome Assembly Stats

The 10X genome assembly resulted in 665,595 contigs contained in 555,643 scaffolds. The total contig and scaffold sequence sizes were 1.704 Gbp and 1.741 Gbp. We believe that this total sequence size is approximately twice the nuclear genome size of around 850 Mbp and that the 10X assembly has largely produced phased very small, phased haplotypes.

The scaffold N50 was 4.1 kbp and the L50 was 99,926 scaffolds. The contig N50 was 3.5 kbp and the L50 was 125,775 contigs. There were 1,242 scaffolds greater than 50 kbp that comprised 5.5% of the genome. There were 189 scaffolds greater than 100 kbp in length, and 2 scaffolds greater than 250 kbp in length.

Overall, this genome would benefit from a long read sequencing technology or another scaffolding technique to increase the N50. Future research plans include determining if the 10X assembly is comprised of phased haplotypes or if it is a pseudohaploid representation of the genome. Due to budgetary constraints, we plan to scaffold the genespace of this genome using RNAseq reads with RAS

#### Transcriptome Assembly Stats

The hybrid transcriptome assembly resulted in 256,027 unique transcripts in 142.1 Mbp. The transcript N50 is 737 bp. There are 30,599 transcripts greater than 1 kb in length, 5,331 transcripts greater than 2.5 kb in length, 689 transcripts greater 5 kb, and 47 transcripts longer than 10 kb.

This transcriptome contains more transcripts than is likely to be biologically relevant, but many of these are likely truncated transcripts from poor input RNA quality from using RNAlater.

#### Peptide mass spectrometry matches

The protein products of all four putative *O. undecimdonta* luciferase transcripts are similar in structure and sequence, except that c9g1i5 lacks 93 N-terminal amino acids. See Figure 7 for a protein alignment and sites where individual peptide peaks were matched to transcripts by the Mascot software.

#### Homologous protein search

The three highest percent identity blastp hits were only 30 amino acids at 40% similarity with and E-value of 0.5 in the Macrochaeta clavicornis transcriptome, 39 amino acids at 35.9% similarity with an E-value of 3.2 in the Harmothoe extenuata transcriptome, and 55 amino acids at 34.55% identity with an E-value of 1.2 in the Chaetopterus variopedatus transcriptome. When these top three hits were used as queries in a blastp search against the NCBI nr database, they had similar or higher percent identities to existing non-luciferase proteins (39.02%, 50%, 60.55%), longer matching regions (41 aa, 50 aa, 218 aa), and varying E-values (2, 3E-16, 7.0E-95). These results generalize to all other blast searches that we conducted. Taken together, these results indicate that there are no conserved proteins in the assembled polychaete transcriptomes given that any blast hit better matches a sequence in the nr database than the protein of transcript c9g1i2.

